# Strict OOD Antigen-to-Antibody Retrieval with CDR-Aware Slot Late Interaction

**DOI:** 10.64898/2026.06.30.735486

**Authors:** Peishuo Liu, Mianzhi Pan, Chenyang Yan, Fuhao Li, Jianbing Zhang

**Affiliations:** National Key Laboratory for Novel Software Technology Nanjing University, Nanjing, Jiangsu, China

**Keywords:** antibody retrieval, antigen-antibody interaction, out-of-distribution evaluation, CDR representation, late interaction, protein language model

## Abstract

Antigen-specific antibody retrieval aims to rank candidate antibodies for a target antigen, providing an early virtual-screening step before structural modeling or experimental validation. Existing sequence-based antibody-antigen interaction studies often formulate the problem as pairwise binding prediction, and random or non-clustered evaluations can over-estimate generalization when related antigens appear across training and test data. We study a strict antigen-cluster out-of-distribution (OOD) retrieval setting in which test antigens come from sequence clusters unseen during training. This setting is difficult because binding is driven by local epitope-CDR complementarity, while available databases mainly contain observed positive complexes and lack reliable negative labels for unlabeled candidates. We propose Ab-CASLR, an antibody CDR-aware slot late-interaction retriever that encodes antigens with ESM-2, encodes antibodies with IgBert, constrains antibody-side latent slots to complementarity-determining regions (CDRs), and scores local slot compatibility instead of single-vector global similarity. On a strict OOD benchmark with 849 antigen queries and 869 candidate antibodies, the model achieves 7.42% Hits@10, outperforming k-mer homology transfer at 5.53% Hits@10 and yielding 6.28-fold enrichment over exact random screening at *K* = 10. Ablations and diagnostics show that CDR-constrained antibody slots remain diverse, whereas antigen-side latent slots collapse into similar summaries. These results support CDR-aware local antibody representation as a useful inductive bias for early binder recovery under strict OOD evaluation, while antigen-side epitope grounding remains unresolved.

## I. Introduction

Antibodies are central to diagnostics, therapeutics, and biological research because they recognize antigens with high specificity [1]. A practical computational problem in antibody discovery is to prioritize candidate antibodies for a target antigen before expensive structural modeling, docking, affinity maturation, or experimental validation [2], [3]. We study antigen-to-antibody retrieval: given an antigen sequence as a query, the model ranks a candidate antibody library so that observed binders are enriched near the top. This formulation matches early virtual screening, where the goal is not to certify all low-ranked antibodies as non-binders but to concentrate plausible binders into a small experimental budget. Accordingly, early-retrieval metrics such as Hits@10 and enrichment factor are more appropriate than aggregate binary accuracy.

This retrieval setting differs from common pairwise binding prediction. A pair classifier receives one antigen-antibody pair and predicts a binding label or score, but it does not directly optimize the within-query ranking of a fixed antibody library. The distinction matters because antibody databases mainly contain observed positive complexes, whereas unpaired antigen-antibody combinations are unlabeled rather than confirmed negatives. A retriever must compare many plausible candidates for the same antigen, exposing ranking failures that independent pair classification can hide.

As illustrated in Fig. 1, antigen-to-antibody retrieval is related to but distinct from pairwise binding classification [4], [5], [6], antibody generation/design [7], [8], [9], and antibody virtual-screening or ranking settings [10], [11]. Rather than predicting a binding label for a single antigen-antibody pair, generating a new antibody sequence, or relying on structure-based screening or affinity ranking, our setting ranks a fixed candidate antibody library for a query antigen using sequence information. This distinction motivates our use of library-level ranking metrics and exact random enrichment baselines.

**Fig. 1.**
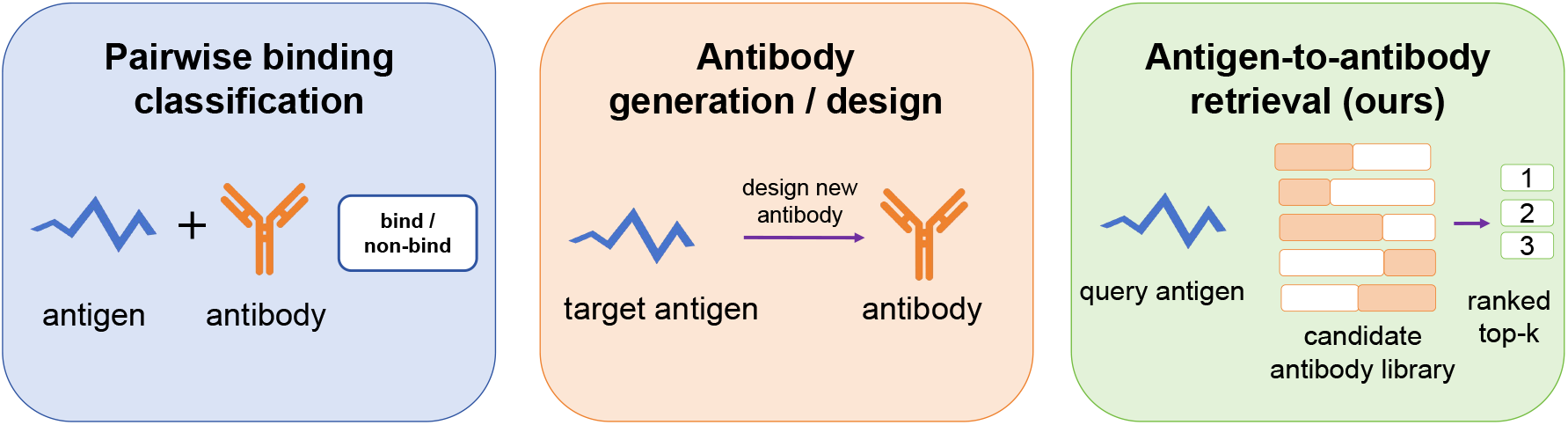
Comparison of three problem settings in computational antibody modeling. Pairwise binding classification predicts a binding score or label for a single antigen-antibody pair. Antibody generation/design aims to design a new antibody for a target antigen. In contrast, antigen-to-antibody retrieval ranks a fixed candidate antibody library for a query antigen, with the goal of enriching observed binders in the top-ranked candidates.

Generalization to unseen antigens is also critical. Random train-test splits can place homologous antigens across splits, allowing models to exploit antigen-family similarity instead of transferable binding signals [12], [13], [14]. We therefore construct a strict antigen-cluster OOD benchmark: antigens are clustered with MMseqs2 at `min_seq_id = 0.8`, and test clusters are excluded from both training and checkpoint selection. The test split contains 849 antigen queries, 869 candidate antibodies, and 872 observed positive pairs. This conservative split asks whether a learned scoring function can retrieve antibodies for antigen sequence families unseen during training.

The biological structure of antigen-antibody recognition motivates local matching. Binding is mediated largely by complementarity between antigen epitopes and antibody CDRs rather than by uniform whole-sequence similarity [15]. Compressing an antibody into a single global embedding can obscure CDR-specific signals. We therefore propose Ab-CASLR, an antibody CDR-aware slot late-interaction retriever: the antibody tower encodes heavy/light chains with IgBert and constructs CDR-constrained slots, the antigen tower encodes the query with ESM-2 and produces latent summaries, and a late-interaction scorer aggregates the strongest local compatibility scores.

On the strict OOD benchmark, the proposed model achieves 7.42% Hits@10 and 6.28-fold enrichment over exact random screening at *K* = 10, outperforming global ESM-2 embedding similarity, k-mer homology transfer, and migrated pair-classification baselines. Ablations and diagnostics show that the gain mainly comes from antibody-side CDR-aware local representation: CDR-constrained antibody slots remain diverse, whereas antigen-side latent slots collapse into highly similar summaries. Thus, the model should be viewed as a retrieval-layer improvement over pretrained sequence representations, not as a validated epitope-paratope decomposition model.

The main contributions of this paper are summarized as follows:

- We introduce a strict antigen-cluster OOD benchmark for antigen-to-antibody library retrieval.
- We propose Ab-CASLR, a CDR-aware slot late-interaction retriever that preserves antibody-side local CDR evidence instead of relying only on global embeddings.
- We report early-enrichment evaluation with exact random baselines, achieving 7.42% Hits@10 and 6.28-fold EF@10 over a controlled corpus of 869 antibodies.
- We provide ablations and diagnostics showing asymmetric slot behavior: antibody slots remain diverse, whereas antigen-side latent summaries collapse.

## II. Related Work

### A. Antigen-Antibody Interaction Prediction

Computational antigen-antibody modeling is often formulated as pairwise interaction or binding prediction, where a model receives one antigen-antibody pair and predicts an interaction label or compatibility score. Sequence-based methods such as AbAgIntPre use antigen and antibody sequence features [4], while recent models incorporate attention, interface-aware modules, pretrained representations, or state-space architectures [5], [16], [6]. These methods are useful for pair scoring but do not directly address antigen-to-antibody library retrieval. Pairwise training also depends on constructed negative pairs, whereas unobserved antigen-antibody combinations are unlabeled. Our work therefore evaluates retrieval directly under a fixed candidate corpus and a strict antigen-cluster OOD split.

### B. Protein and Antibody Language Models

Protein language models provide transferable sequence representations for biological tasks. General encoders such as ESM-style models capture evolutionary and structural regularities [17], while antibody-specific language models such as AntiBERTa, AbLang, AntiBERTy, IgBert, and IgT5 better reflect immunoglobulin sequence distributions [18], [19], [20], [21]. A direct retrieval baseline is to pool antigen and antibody representations into global embeddings. However, antigen-antibody recognition is asymmetric and local: antigen sequences may contain regions unrelated to binding, whereas antibody specificity is concentrated in CDRs [15]. Our model keeps pretrained sequence encoders but replaces global embedding similarity with CDR-aware local matching.

### C. OOD Evaluation and Data Leakage

Evaluation protocol is critical in biological sequence modeling. Random splits can place homologous proteins across training and test sets, leading to optimistic estimates of generalization [12], [13], [14]. These studies motivate cluster-based splits for evaluating transfer to unseen sequence families. Following this principle, test antigen clusters in our benchmark are excluded from both training and checkpoint selection.

### D. Late Interaction and Local Matching

Late-interaction retrieval models preserve multiple local representations and compute relevance through token- or region-level matching instead of single-vector similarity. ColBERT retains contextualized token embeddings and performs late interaction at scoring time [22]. We adapt this idea by constructing CDR-constrained antibody slots and matching them with antigen-side latent summaries through max-compatible local scoring.

## III. Strict ood Antigen-to-Antibody Retrieval Benchmark

### A. Task Definition

Given an antigen query *q* and a fixed candidate antibody corpus *D*, the task is to rank antibodies *d* ∈ *D* such that observed binders appear near the top. Each query may have one or more observed positive antibodies. Because unobserved antigen-antibody pairs are unlabeled rather than reliable negatives, we evaluate top-*K* recovery and enrichment instead of binary accuracy over all unpaired candidates.

Formally, the model learns a scoring function *f* (*q, d*). For each query *q*, all candidates in *D* are sorted in decreasing order of *f* (*q, d*). Let *P*_*q*_ ⊆ *D* denote the set of observed positive antibodies for *q*, and let *n*_*q*_ =|*P* _*q*_ |. The induced ranking is written as

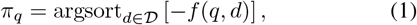

where *π*_*q*_(*r*) denotes the antibody ranked at position *r*. The binary relevance label is defined as

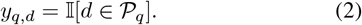

Reported scores measure recovery of observed positives rather than exhaustive identification of all possible binders.

Let *Q* denote the set of antigen queries and *N* = |*D*|denote the candidate corpus size. Hits@*K* measures the fraction of queries for which at least one observed positive antibody is retrieved in the top *K*:

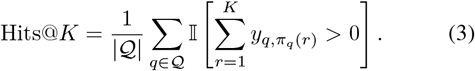

Recall@*K* measures the fraction of observed positive pairs covered by the top-*K* ranked candidates:

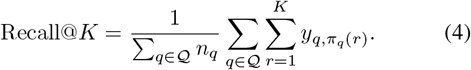

We also report MRR@10 and NDCG@10.

To quantify screening value relative to random selection, we compute an exact random baseline for each query. For a query with *n*_*q*_ observed positives in a corpus of *N* candidates, the probability that random sampling without replacement retrieves at least one positive in *K* trials is 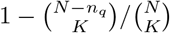.

Thus,

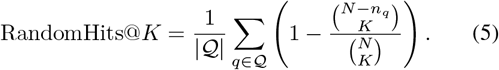

The enrichment factor is then defined as

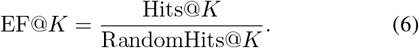

### B. Data Source and Pair Construction

We construct the benchmark from SAbDab-derived antigen-antibody structural complexes [23]. Each record is converted into a retrieval pair in which the antigen sequence is the query and the paired antibody is an observed positive candidate. We retain antigen entries with non-empty amino-acid sequences, yielding a protein-sequence antigen benchmark. Multi-chain antigens are represented by concatenating chain sequences with a delimiter, and antibodies are represented by available heavy- and light-chain sequences.

Duplicate structures are handled at the sequence-pair level rather than only by structure identifier. Specifically, records are grouped by antigen sequence, binder type, heavy-chain sequence, and light-chain sequence. Within each duplicate group, an anchor record is selected and other records are compared with the anchor using contact-derived epitope and paratope labels. Singleton, anchor, and contact-distinct records are retained; contact-duplicate and ambiguous records are removed.

We do not construct confirmed negatives from unpaired antigens and antibodies, because the absence of an observed complex does not imply lack of binding. For each antigen query, observed binding antibodies are positives, while other candidates are unlabeled during ranking evaluation.

### C. Antigen-Cluster OOD Split

Antigen sequences are clustered using MMseqs2 [24] with `min_seq_id = 0.8.` All antigens within the same sequence cluster are assigned to the same split, so no antigen cluster appears in more than one of the training, validation, and test sets. This design reduces homologous antigen leakage and evaluates whether a model can retrieve antibodies for antigens from sequence families that are unseen during training.

Table I summarizes the split statistics. The split is stricter than a pair-level random split because closely related antigens cannot contribute examples to both optimization and evaluation. The resulting test set contains 849 antigen queries, 869 unique candidate antibodies, and 872 observed positive antigen-antibody pairs. Thus, each method must rank hundreds of candidate antibodies for unseen antigen queries.

**TABLE I.** Strict ood benchmark statistics. The test antibody set is the controlled candidate corpus.

| Split | Pairs | Antigens | Antibodies |
| --- | --- | --- | --- |
| Train | 5698 | 5258 | 5692 |
| Validation | 818 | 799 | 818 |
| Test | 872 | 849 | 869 |

In the test split, 21 antigen queries have more than one observed positive antibody. Test antigen clusters are not used for training or checkpoint selection.

### D. Candidate Corpus and Evaluation Protocol

The test candidate corpus consists of the 869 unique antibodies appearing in the test split. For every test antigen query, all methods rank this same corpus, and an antibody is counted as positive only if the corresponding antigen-antibody pair is observed in the curated benchmark. All other combinations are treated as unlabeled candidates. This controlled corpus avoids comparing methods under different retrieval spaces and makes exact random enrichment computable. It should be interpreted as a benchmark corpus rather than a deployment-scale antibody library. Because absolute Hits@*K* depends on corpus size and the number of observed positives per query, we report both raw top-*K* recovery and EF@*K* over exact random selection.

## IV. Ab-CASLR: Antibody cdr-Aware Slot Late-Interaction Retriever

### A. Overview

The model is a heterogeneous dual-tower retriever. The query tower encodes antigen sequences with ESM-2 150M, and the document tower encodes available antibody heavy- and/or light-chain sequences with IgBert. Both pretrained encoders are fine-tuned with a small learning rate, and their token-level representations are projected into a shared *h*-dimensional retrieval space with *h* = 128. For an antigen query *q* and an antibody candidate *d*, we denote the projected token representations as

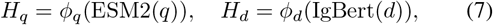

where 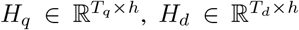, and *ϕ*_*q*_, *ϕ*_*d*_ are learned linear projections.

Instead of compressing each antibody into a single global embedding, the document tower constructs multiple CDR-aware latent slots. These slots are constrained by CDR masks and aggregate information from the H1, H2, H3, L1, L2, and L3 regions when the corresponding chains are available. This design introduces an antibody-side local inductive bias: antibody specificity is expected to be concentrated in CDR loops, whereas non-CDR regions are not used as primary slot evidence. The two towers are intentionally asymmetric: the antigen tower summarizes a general protein sequence without predefined epitope annotations, while the antibody tower uses CDR annotations as biologically meaningful local regions.

The antigen tower produces *L* = 8 latent query summaries, and the antibody tower produces *M* = 8 CDR-aware document slots. A low-rank bilinear compatibility function compares query summaries with antibody slots. The final retrieval score is computed by max-compatible late interaction, allowing each antigen summary to match its most compatible antibody CDR slot. Figure 2 shows the model overview.

**Fig. 2.**
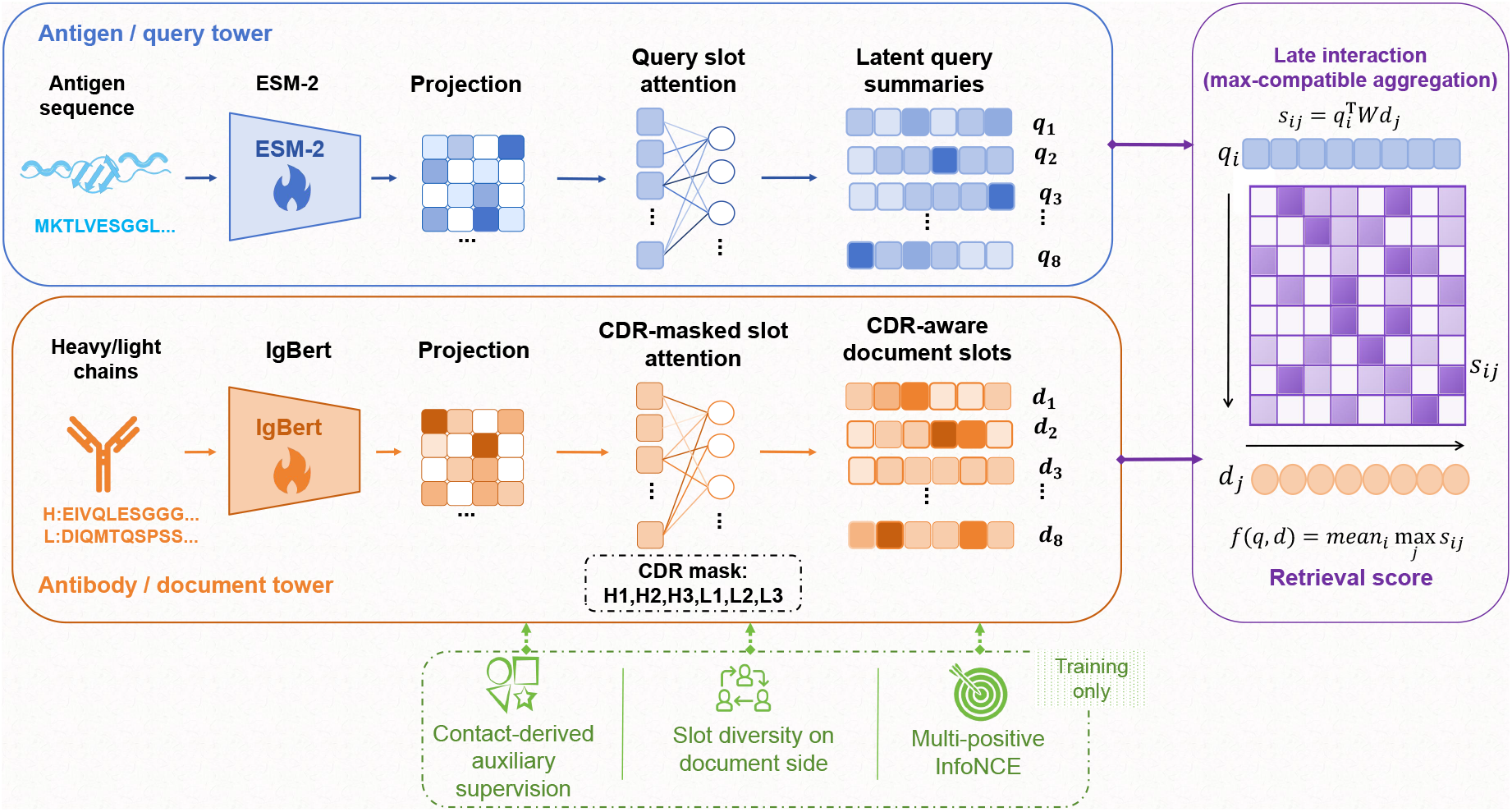
Overview of the CDR-aware slot late-interaction retriever. ESM-2 produces antigen query summaries, IgBert produces CDR-constrained antibody slots, and a low-rank bilinear late-interaction layer aggregates local compatibility scores. Auxiliary contact and diversity losses are used only during training.

### B. Antibody-Side CDR Slot Construction

Let 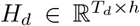 denote the projected IgBert token representations of an antibody candidate. We construct *M* = 8 document slots on the antibody side. Unlike global pooling, slot attention is constrained by CDR masks so that document slots aggregate CDR information. Let *c*_*j*_ ∈ {0, 1}indicate whether antibody token *j* belongs to an annotated CDR region. For slot *r*, the unnormalized attention logit for token *j* is denoted as *a*_*rj*_. The CDR-masked attention weights are defined as

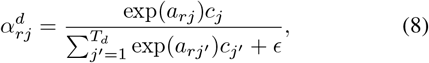

and the resulting antibody slot is

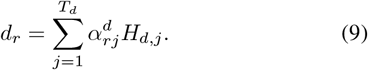

This construction introduces an antibody-side local inductive bias. The CDR mask restricts primary slot evidence to CDR tokens, while the pretrained antibody encoder still contextualizes each token within the available sequence. The CDR masks are defined according to the Chothia numbering scheme [25]. When both heavy and light chains are available, the mask covers H1, H2, H3, L1, L2, and L3 regions. For entries with a single available chain, only the corresponding CDR regions are used.

The resulting slots provide multiple local antibody representations for late interaction, giving the scoring layer CDR-level evidence instead of a single pooled antibody vector. A document-side diversity regularizer reduces redundant slots during training.

### C. Antigen-Side Latent Summaries

The antigen query tower uses the projected ESM-2 token representations 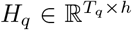 to construct *L* = 8 latent query summaries. Unlike the antibody side, the antigen side has no predefined CDR-like regions, so we use unconstrained slot attention to aggregate antigen token features. For query summary *i*, the attention weight over antigen token *j* is

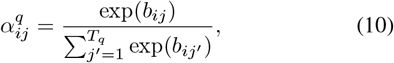

and the corresponding latent summary is

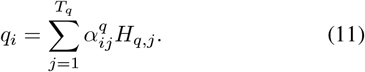

These summaries are learned antigen-side retrieval features and are not assumed to correspond directly to experimentally validated epitopes. Their empirical behavior is analyzed in the slot-space diagnostics.

### D. Late-Interaction Scoring

Given antigen query summaries 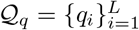and antibody CDR slots 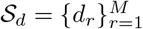, we compute local antigen-antibody compatibility with a low-rank bilinear scorer. For query summary *q*_*i*_ and antibody slot *d*_*r*_, the compatibility score is

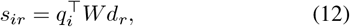

where *W* ∈ ℝ^*h×h*^ is parameterized as *W* = *UV* ^⊤^, with *U, V* ∈ ℝ^*h×R*^ and rank *R* = 64.

The final retrieval score keeps the maximum compatibility over antibody slots for each query summary and averages these maxima:

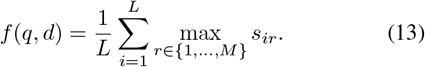

This late interaction is less restrictive than global embedding similarity and remains suitable for library retrieval because antibody candidates can be encoded into reusable local slots before ranking.

### E. Training Objective

The model is trained with a multi-positive contrastive retrieval objective in which all observed positives for a query appearing in the same batch are valid positives. In addition to the retrieval loss, we use contact-derived epitope/paratope auxiliary supervision and document-side slot diversity regularization.

For the retrieval objective, let ℬ denote a training batch, *Q*_*ℬ*_ denote the antigen queries in the batch, and *D*_*ℬ*_ denote the antibody candidates used as in-batch retrieval candidates. For query *q*, let 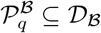 denote the set of observed positive antibodies for *q* that appear in the batch. The multi-positive contrastive loss for query *q* is

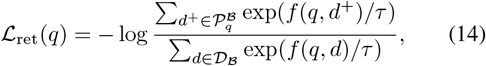

where *τ* is the contrastive temperature. The batch retrieval loss is averaged over queries with at least one in-batch positive:

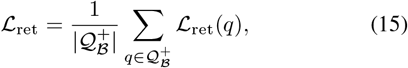

where ^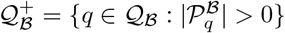^.

We further use structural contact labels as auxiliary supervision when available. For structurally supervised positive pairs, antigen and antibody residues are marked as contacts when their minimum all-atom distance is at most 4.0 *Å*. Rows without contact annotations remain valid retrieval pairs but do not contribute to the auxiliary losses. Let 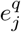 and 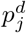 denote binary residue-level contact labels for antigen token *j* and antibody token *j*, respectively. Missing labels are ignored by validity masks.

The epitope and paratope auxiliary losses encourage slot-attention coverage to align with these residue-level contact labels. Let Ω_*q*_ and Ω_*d*_ denote the sets of valid labeled antigen and antibody tokens. We define

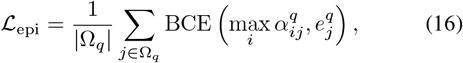

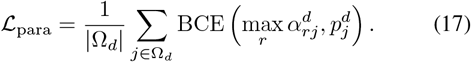

Here, max-pooled attention measures whether a token is covered by any query summary or document slot.

To reduce redundant antibody-side slots, we penalize pairwise cosine similarity among document slots:

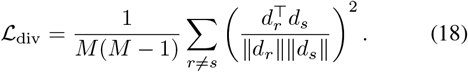

The full training objective is

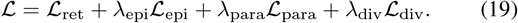

The auxiliary terms are used as regularizers for slot learning rather than as standalone evidence of mechanistic correctness; the final evaluation is based on strict OOD retrieval ranking.

## V. Experiments

### A. Evaluation Metrics

We report Hits@*K*, Recall@*K*, MRR@10, NDCG@10, and EF@*K* following Section III. Hits@10 and EF@10 are primary because the target use case is early candidate prioritization for downstream modeling or experimental validation. MRR@10 and NDCG@10 measure the rank position of recovered positives; Hits@50 and Hits@100 reflect broader shortlist quality. Because only 21 test queries contain multiple observed positive antibodies, Hits@*K* and EF@*K* are the most interpretable metrics for this benchmark.

### B. Implementation Details

We fine-tune ESM-2 150M and IgBert with learning rate 3 × 10^*−*6^, and train the newly introduced projection, slot-attention, and compatibility modules with learning rate 1 × 10^*−*4^. The shared retrieval dimension is 128, with *L* = 8 query summaries, *M* = 8 document slots, and bilinear rank 64.

The model is trained for 30 epochs with a batch size of 8. The loss weights are set to *λ*_epi_ = 0.01, *λ*_para_ = 0.01, and *λ*_div_ = 0.001. Model selection is performed on the validation split, and the selected checkpoint is evaluated once on the held-out strict OOD test split. All methods are evaluated on the same 849 test antigen queries and the same 869-antibody candidate corpus.

### C. Main Results and Random Enrichment

Table II reports the retrieval and enrichment results of the validation-selected checkpoint on the strict OOD test split. The proposed retriever achieves 7.420% Hits@10. Although the absolute top-*K* recovery remains modest, this result should be interpreted relative to the controlled candidate corpus and the exact random baseline. With 869 candidate antibodies per query, exact random selection obtains 1.182% Hits@10. Our model therefore yields a 6.28-fold enrichment over random screening at *K* = 10.

**TABLE II.** Strict ood retrieval and enrichment results.

| $K$ | Random Hits@$K$ | Model Hits@$K$ | EF@$K$ |
| --- | --- | --- | --- |
| 1 | 0.118% | 1.767% | 14.97× |
| 5 | 0.591% | 4.594% | 7.77× |
| 10 | 1.182% | **7.420%** | **6.28×** |
| 50 | 5.900% | 16.726% | 2.83× |
| 100 | 11.781% | 25.206% | 2.14× |

The enrichment is strongest at small *K*. At *K* = 1, the model achieves 1.767% Hits@1 and 14.97×EF@1, showing strong enrichment but not enough reliability for standalone recommendation. At *K* = 10, the model maintains a practical shortlist size for downstream modeling or experimental triage. As *K* increases, EF@*K* decreases because random screening has a higher probability of recovering at least one observed positive.

At *K* = 10, the model obtains 7.362% Recall@10, 2.936% MRR@10, and 3.831% NDCG@10, consistent with enrichment near the top of the ranked list.

### D. Comparison with External Baselines

Table III compares the proposed retriever with external baselines under the same strict OOD test corpus. We include k-mer homology transfer, global ESM2-ESM2 embedding similarity, and migrated pair-classification models: DeepInterAware [16], MambaAAI [6], and AntiBinder [5]. For k-mer homology transfer, each test antigen first retrieves the top five most similar training antigens using 4-mer Jaccard similarity. It then transfers the antibodies paired with these training antigens to the test antibody corpus using 4-mer Jaccard similarity over concatenated heavy/light antibody sequences. Let *K*_4_(*x*) denote the set of 4-mers extracted from sequence *x*. The sequence similarity between two sequences is defined as

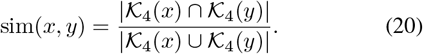

**TABLE III.** External baseline comparison under the same strict ood test corpus. Values are hits@*k*.

| Method | Hits@1 | Hits@10 | Hits@50 |
| --- | --- | --- | --- |
| k-mer homology transfer | 1.65% | 5.53% | 14.84% |
| ESM2-ESM2 embedding similarity | 1.29% | 3.29% | 8.12% |
| DeepInterAware (1v1) | 0.00% | 1.83% | 7.45% |
| DeepInterAware (1v5) | 0.57% | 4.36% | 11.70% |
| DeepInterAware (1v10) | 0.23% | 2.87% | 12.96% |
| MambaAAI (1v1) | 0.24% | 2.00% | 4.95% |
| MambaAAI (1v5) | 0.00% | 1.88% | 3.89% |
| AntiBinder (1v1) | 0.24% | 1.70% | 11.63% |
| AntiBinder (1v5) | 0.00% | 1.50% | 7.40% |
| **Ab-CASLR** | **1.77%** | **7.42%** | **16.73%** |

For a test antigen *q* and a candidate antibody *d*, the homology-transfer score is

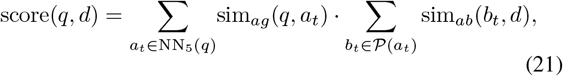

where NN_5_(*q*) denotes the top five training antigens most similar to *q*, and *P*(*a*_*t*_) denotes the training antibodies observed to bind antigen *a*_*t*_. For antibody similarity, heavy and light chains are concatenated before 4-mer extraction. All methods use the same 849 test antigen queries and the same 869-antibody candidate corpus. Each method scores every antigen-candidate pair and ranks candidates by the resulting score. For pair-classification baselines, the classifier output is directly used as the retrieval score. The suffixes 1v1, 1v5, and 1v10 denote the positive-to-sampled-negative ratio used when adapting the pairwise classifiers; they do not change the candidate corpus at evaluation time. Thus, no baseline is evaluated on a smaller or easier retrieval space. All migrated pair-classification baselines are trained on the same training split and selected using the same validation split before test-corpus ranking.

Ab-CASLR achieves the highest Hits@10 among all tested baselines. Compared with k-mer homology transfer, Hits@10 improves from 5.53% to 7.42%, an absolute gain of 1.89 percentage points and a relative improvement of 34.2%. Compared with global ESM2-ESM2 embedding similarity, our model more than doubles Hits@10, indicating that global sequence or embedding similarity is insufficient for strict OOD antigen-to-antibody retrieval. Migrated pair-classification baselines perform worse, suggesting that scores learned under pairwise negative construction do not directly transfer to within-query library ranking.

### E. Component Ablations

Table IV reports component ablations under the same strict OOD split, the same 869-antibody candidate corpus, and the same validation-based checkpoint selection rule. These runs are diagnostic rather than fully matched one-factor hyperparameter sweeps, but they test the main architectural choices of the proposed retriever.

**TABLE IV.** Component ablations under the strict ood test protocol. Values are test hits@10.

| Variant | Hits@10 | Main change |
| --- | --- | --- |
| Full model | **7.420%** | Proposed retriever |
| Global pooled dual-tower | 1.413% | Remove slots |
| w/o CDR mask | 6.360% | Full-sequence slots |
| CDR pooled similarity | 2.120% | Remove late interaction |
| Single query summary | 5.065% | Reduce query slots |
| Mean aggregation | 4.594% | Replace max aggregation |

The results support CDR-aware local matching. A global pooled dual-tower model achieves only 1.413% Hits@10. Removing the CDR mask reduces Hits@10 to 6.360%, and replacing late interaction with pooled CDR similarity reduces it to 2.120%. Single-query and mean-aggregation variants also remain below the full model, indicating that the full multi-summary and max-aggregation configuration is empirically preferable, although the diagnostics below show that the learned antigen-side summaries are not diverse.

### F. Slot-Space Diagnostics

We further analyze learned slot representations to understand the retrieval gain. Table V reports mean off-diagonal cosine similarity among raw slots, compatibility-projected slots, slot-attention distributions, and mean attention entropy. The diagnostics reveal a clear asymmetry between the antigen and antibody sides. On the antigen side, query slots are nearly identical: the raw slot cosine, compatibility-projected slot cosine, and attention pairwise cosine are all 1.000 after rounding. Thus, learned query summaries should not be interpreted as distinct epitope-like regions. On the antibody side, the document slots are substantially more diverse. The raw slot cosine is 0.465, and the attention pairwise cosine is only 0.107, showing that different antibody slots attend to different CDR-level patterns. The lower antibody-side entropy is consistent with more localized attention, although entropy is length-dependent and should be treated as supporting evidence. These results clarify the mechanism. Although the architecture uses slots on both sides, the empirical behavior does not support a claim of validated joint epitope-paratope decomposition. The main effective mechanism is CDR-aware antibody-side local representation combined with late interaction; reliable antigen-side epitope grounding remains unresolved.

**TABLE V.**
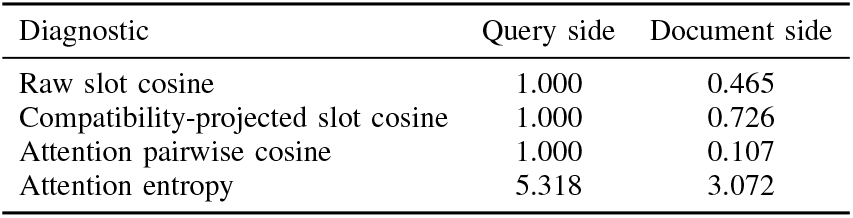
Slot-space diagnostics of the validation-selected full model. Values are means over slot pairs or attention distributions; cosine values are off-diagonal means.

## VI. Discussion and Conclusion

We presented a strict antigen-cluster OOD benchmark and a CDR-aware slot late-interaction retriever for antigen-to-antibody retrieval. Under a controlled test setting with 849 antigen queries and 869 candidate antibodies, Ab-CASLR achieves 7.420% Hits@10 and 6.28-fold enrichment over exact random screening. It also outperforms k-mer homology transfer, global ESM2 embedding similarity, and migrated pair-classification baselines under the same candidate corpus. Ablations and diagnostics indicate that the main benefit comes from antibody-side CDR-aware local representation combined with max-compatible late interaction. Global pooled matching performs poorly, removing CDR constraints weakens retrieval, and replacing late interaction with pooled CDR similarity substantially reduces Hits@10. At the same time, antigen-side latent summaries collapse into highly similar representations, so the current model should not be interpreted as a validated epitope-paratope decomposition model.

The study has limitations. The benchmark contains observed positive complexes but no experimentally verified negatives for all unpaired candidates. The candidate corpus is a controlled test-set antibody library rather than a deployment-scale screening database. The model is sequence-centered and does not explicitly evaluate three-dimensional loop conformations, antigen surface exposure, binding affinity, or induced fit. Future work should improve antigen-side binding-region grounding with structure-aware antigen representations, surface accessibility features, residue-level epitope supervision, or curated negative evidence from binding assays. Overall, these findings support CDR-aware local retrieval for early antibody prioritization under strict OOD evaluation while identifying antigen-side epitope grounding as a central bottleneck. The reported results are based on a single strict OOD split, and future work should evaluate robustness across additional antigen-cluster splits and larger candidate libraries.

